# Detection of race-specific resistance against *Puccinia coronata* f. sp. *avenae* in *Brachypodium* species

**DOI:** 10.1101/281568

**Authors:** Vahid Omidvar, Sheshanka Dugyala, Feng Li, Susan Rottschaefer, Marisa E. Miller, Mick Ayliffe, Matthew J. Moscou, Shahryar F. Kianian, Melania Figueroa

## Abstract

Oat crown rust caused by *Puccinia coronata* f. sp. *avenae* is the most destructive foliar disease of cultivated oat. Characterization of genetic factors controlling resistance responses to *Puccinia coronata* f. sp. *avenae* in non-host species could provide new resources for developing disease protection strategies in oat. We examined symptom development and fungal colonization levels of a collection of *Brachypodium distachyon* and *B. hybridum* accessions infected with three North American *P. coronata* f. sp. *avenae* isolates. Our results demonstrated that colonization phenotypes are dependent on both host and pathogen genotypes, indicating a role for race-specific responses in these interactions. These responses were independent of the accumulation of reactive oxygen species. Expression analysis of several defense-related genes suggested that salicylic acid and ethylene-mediated signaling, but not jasmonic acid are components of resistance reaction to *P. coronata* f. sp*. avenae*. Our findings provide the basis to conduct a genetic inheritance study to examine if effector-triggered immunity contributes to non-host resistance to *P. coronata* f. sp. *avenae* in *Brachypodium* species.

## Introduction

Oat crown rust caused by the obligate biotrophic rust fungus *Puccinia coronata* f. sp. *avenae* is the most widespread and damaging foliar disease of cultivated oat (*Avena sativa* L.) (Nazareno et al. 2017). Symptoms of crown rust infection manifest in the foliar tissue, causing a reduction in photosynthetic capacity and thus affecting grain size and quality (Holland and Munkvold 2001; Simons 1985). The most sustainable method to control this devastating disease is the use of genetic resistance (Carson 2011). Novel sources of genetic resistance may therefore translate into novel crop protection strategies.

In general, plant disease resistance to would-be pathogens can be conferred by either constitutive or induced barriers (Heath 2000; Nurnberger and Lipka 2005). Physical barriers, such as rigid cell walls and waxy cuticles as well as preformed antimicrobial compounds are some of the constitutive obstacles that explain why plants are immune to most microbes. Nevertheless, certain microbes have evolved strategies to overcome these barriers, and in such instances plant-microbe incompatibility is based upon pathogen recognition and the induction of plant defense responses (Heath 2000, Periyannan et al. 2017).

Inducible defense responses in plants are mediated by a tightly regulated two-tier immune recognition system that, depending on the physiological characteristics of the potential pathogen, may or may not be effective at preventing microbial colonization (Dodds and Rathjen 2010). The first layer of microbial recognition is controlled by cell surface associated receptors, named pattern recognition receptors (PRRs). PRRs recognize conserved and essential microbial molecules known as pathogen-associated molecular patterns (PAMPs) and/or plant derived damage-associated molecular patterns (DAMPs). This recognition leads to the activation of a range of broad-spectrum basal defenses that constitute PAMP-triggered immunity (PTI) (Zipfel 2008). PAMPs in pathogenic fungi like the crown rust pathogen include chitin or xylanases, which are essential constituents of the fungal cell wall. In contrast, DAMPs are not necessarily pathogen derived and include plant cell wall fragments and plant peptides released during infection.

Adapted pathogens manipulate PTI signaling events and suppress basal defenses by secreting a suite of effector proteins into plant cells, thereby enabling successful plant colonization in some instances (Toruno et al. 2016). However, disease resistance against adapted pathogens can still occur if the plant can recognize one or more of these secreted effector molecules. Effector recognition generally occurs by plant intracellular nucleotide binding leucine rich repeat (NB-LRR) receptors that each induce defense responses upon recognition of a cognate pathogen effector, also known as an avirulence (Avr) protein. This second layer of pathogen recognition, referred to as effector-triggered immunity (ETI), frequently results in a localized hypersensitive cell death response at attempted infection sites (HR) and is the underlying molecular basis of the gene-for-gene concept (Dodds and Rathjen 2010; Ellis et al. 2014; Flor 1971). ETI is highly specific with resistance only occurring if the adapted pathogen isolate expresses an effector that is recognized by a corresponding NB-LRR in the infected plant.

The delivery of rust effector proteins into host plants is mediated by distinct fungal infection structures called haustoria (Catanzariti et al. 2006; Garnica et al. 2014; Panstruga and Dodds 2009). Only a small number of effectors have been characterized in rust fungi (Chen et al. 2017; Maia et al. 2017; Ravensdale et al. 2011; Salcedo et al. 2017), however, genome-wide effector mining suggests that rust pathogens may deploy hundreds of effector molecules via the haustorium during infection (Cantu et al. 2013; Duplessis et al. 2011; Hacquard et al. 2012; Nemri et al. 2014; Rutter et al. 2017). Oat crown rust populations are typified by a complex race-structure, which likely originates from variation in the array of effectors present in different pathogen genotypes (Carson 2011; Chong et al. 2000; Nazareno et al. 2017). For years, oat breeding programs have relied upon naturally occurring resistance in *Avena* spp., most likely mediated by *NB-LRR* genes, to protect against crown rust. However, the resistance of these oat varieties is often rapidly overcome by evolution of the crown rust pathogen to avoid host plant recognition. Achieving increased durability in oat crown rust resistance therefore requires new sources of genetic immunity to be identified and further advances made in our understanding of the molecular basis of rust recognition in host and non-host plant species.

Non-host plant species potentially offer an untapped resistance resource. Non-host resistance (NHR) is typically described in the context of a dichotomy that distinguishes it from host resistance. NHR is defined as genotype-independent and effective against all genetic variants of a non-adapted pathogen species, whereas host resistance is genotype-dependent and effective only against a subset of genetic variants of an adapted pathogen (Mysore and Ryu 2004). This paradigm has been gradually changing as accumulating evidence suggests that signaling pathways and defense mechanisms overlap in non-host and host resistance and that microbial adaptation to plant species does not conform to a simple qualitative distinction (Bettgenhaeuser et al. 2014; Dawson et al. 2015; Figueroa et al. 2015; Gill et al. 2015; Thordal-Christensen 2003). Instead, disease phenotypes observed for different plant and pathogen interactions span a continuous spectrum of outcomes ranging from immunity to susceptibility, with many intermediate interactions, making the classification of non-host versus host systems problematic (Bettgenhaeuser et al. 2014; Dawson et al. 2015). Regardless of the terminology, identifying the molecular determinants of non-host or intermediate resistance is of great interest as this type of resistance could be durable and broad-spectrum and contribute significantly to crop improvement.

*Brachypodium distachyon*, a small grass closely related to cereals, is considered a non-host to several rust fungi species related to *P. coronata* f. sp. *avenae*, such as *Puccinia emaculata, Puccinia striiformis, Puccinia graminis* and *Puccinia triticina* (Ayliffe et al. 2013; Barbieri et al. 2011; Bettgenhaeuser et al. 2014; Bossolini et al. 2007; Dawson et al. 2015; Figueroa et al. 2013, 2015; Gill et al. 2015). Variation in fungal colonization in some of these interactions suggests that it may be possible to genetically dissect these types of immune responses and harness them for engineering rust disease resistance (Figueroa et al. 2015). In this study, we examined the interaction between *P. coronata* f. sp. *avenae* and *B. distachyon*, as well as *Brachypodium hybridum. B. hybridum* is an allotetraploid species originated from the hybridization of the diploid species *B. distachyon and B. stacei* (Lopez-Alvarez et al. 2012).

Pathogen susceptibility in plants, particularly to rust fungi, is a process, which remains poorly characterized. Our study shows that *P. coronata* f. sp. *avenae* can infect *B. distachyon* and *B. hybridum* leaves and grow extensively, but cannot sporulate. These findings open the possibility of investigating mechanisms that confer partial rust susceptibility in *Brachypodium*. Gene expression analysis of an ortholog of the putative rust susceptibility factor in wheat, which encodes a hexose transporter (Moore et al. 2015), suggested that sugar transport may also be important to sustain the growth of *P. coronata* f. sp. *avenae* in *B. distachyon*. In addition to this, we found evidence for race-specific resistance in both species supporting the model proposed by Schulze-Lefert and Panstruga (2011), which postulates a role of ETI in NHR due to the phylogenetic relatedness between the non-host and host species. These findings provide a framework to conduct genetic inheritance studies to dissect recognition of *P. coronata* f. sp. *avenae* by *B. distachyon* and *B. hybridum* and identify loci governing intermediate oat crown rust susceptibility.

## Materials and Methods

### Plant and fungal materials

Twenty-two accessions of *B. distachyon* (ABR6, ABR7, Adi12, Adi13, Adi15, Bd1-1, Bd18-1, Bd21, Bd21-3, Bd2-3, Bd29-1, Bd30-1, Bd3-1, BdTR10H, BdTR13K, Foz1, J6.2, Jer1, Koz5, Luc1, Mon3, and Tek4) and three accessions of *B. hybridum* (Bou1, Bel1, and Pob1) were used in this study. Seeds were obtained from the John Innes Centre (Dawson et al. 2015), Aberystwyth University (Mur et al. 2011), Joint Genome Institute and Montana State University (Vogel et al. 2009), Universidad Politécnica de Madrid (Dr. Elena Benavente) and USDA-ARS Plant Science Unit, St. Paul, MN, U.S.A (Garvin et al. 2008). All plants were increased by single seed descent prior to conducting this study. The cultivated oat (*Avena sativa*) variety Marvelous was used as a susceptible host to *P. coronata* f. sp. *avenae*. This study used three North American oat crown rust isolates, 12NC29 (race LBBB) and 12SD80 (race STTG) (Miller et al. 2018) collected in 2012, and a historic race 203 (race QBQT) known to be avirulent to the oat variety Victoria (Chang and Sadanaga 1964). Isolates were obtained from the rust collection available at the USDA-ARS Cereal Disease Laboratory, St. Paul, MN, U.S.A., and physiological race analysis was conducted following a standard nomenclature system using a set of oat differentials (Chong et al. 2000; Nazareno et al. 2017). Reported race assignment conveys consistent results from three independent experiments. Virulence phenotypes of *P. coronata* f. sp. *avenae* isolates on oat differentials with infection types of “0”, “0;”, “;”, “;C”, “1;”, “1”, “2”, “3”, “3+”, and “4” were converted to a 0-9 numeric scale, respectively for heat map generation.

### Inoculation and pathogen assays

*Brachypodium* and oat seedlings were grown with 18/6 h light/dark, 24/18°C day/night cycles and 50% humidity. Urediniospores were activated by heat shock treatment at 45°C for 15 min to break cold induced dormancy and suspended in an oil carrier (Isopar M, ExxonMobil) for spray inoculation. Seedlings of *Brachypodium* accessions and oat were inoculated using 50 μL of each *P. coronata* f. sp. *avenae* inoculum (10 mg urediniospores/mL) at three-leaf and first-leaf stages, respectively. For mock inoculation, seedlings were sprayed with oil without urediniospores. Infected seedlings were placed in dew chambers in the dark for 12 h with intermittent misting for 2 min every 30 min. After 12 h, misting was stopped and seedlings were exposed to light for 2 h before they were placed back in growth chambers. Infected primary leaves of oat and secondary leaves of *Brachypodium* accessions were collected for analysis. Presence (+) or absence (-) of macroscopic symptoms, including chlorosis and/or necrosis as well as severity of the symptoms (shown as increments of +) were evaluated at 14 days post infection (dpi) in two independent experiments (biological replicates). Each biological replicate simultaneously tested all accessions, and five plants were examined per accession. Each plant was considered a technical replicate within one independent experiment. Digital images were captured using a stereomicroscope (Olympus model SZX16). To examine correlations between necrosis or chlorosis and estimates of fungal colonization (mean values), symptoms described as “−”, “+”, “++”, “+++” and “++++” were transformed to numerical values 0-4, respectively, to calculate a Spearman’s rank correlation coefficient.

### Analysis of fungal colonization by microscopy

Both infected secondary *Brachypodium* leaves at 14 dpi and infected primary *A. sativa* (variety Marvelous) leaves at 12 dpi were cut into 1 cm length sections and stained with wheat germ agglutinin Alexa Fluor^®^ 488 conjugate (WGA-FITC; ThermoFisher Scientific) to visualize fungal colonies as previously described (Ayliffe et al. 2013; Dawson et al. 2015). Visualization of fungal intracellular growth was carried out using a fluorescence microscope (Leica model DMLB) under blue light with a 450-490 nM excitation filter. The percentage of leaf colonized (pCOL) by *P. coronata* f. sp. *avenae* was estimated according to the method of Dawson et al. (2015) with a modification to report 0, 0.5, 0.75, and 1 scores for disjointed fields of view with hyphal growth less than 15%, between 15-50%, 50-75%, and greater than 75%, respectively. Two independent experiments (biological replicates) were evaluated per *P. coronata* f. sp. *avenae* isolate (12SD80, 203, and 12NC29) and each biological replicate included three leaves. Each leaf was considered a technical replicate within one independent experiment. pCOL values from all three leaves were combined to obtain a mean and standard error of the mean. Fungal development was also examined in whole-mounted infected leaves of *B. distachyon* accessions ABR6 and Bd21 and oat at 1, 2, 3, and 6 dpi. The percentage of urediniospores that successfully germinated and formation of various infection structures, including appresorium (AP), substomatal vesicle (SV), haustorium-mother cell (HMC), and established colonies (EC), were recorded in WGA-FITC stained samples. Three independent experiments (biological replicates) were conducted and simultaneously evaluated infections with *P. coronata* f. sp. *avenae* isolates 12SD80, 203, and 12NC29. Each biological replicate recorded fungal development for 100 infection sites. Mean and standard error of the mean per infection structure category were calculated based on all three biological replicates.

### Analysis of H_2_O_2_ accumulation

H_2_O_2_ accumulation was evaluated using 3,3’-diaminobenzidine (DAB) staining as described by Thordal-Christensen et al. (1997). Infected leaves were stained in 1 mg/mL DAB aqueous solution at pH 3.8 for 4 h in dark and destained in Farmer’s fixative for 12 h. Three independent experiments (biological replicates) were conducted. Each biological replicate included one to two infected secondary leaves to record the number of urediniospores from a 100 sample within a 500 μm distance to a DAB staining site at 2, 4 and 6 dpi. The oat differential that contains the resistance gene *Pc91*, which confers resistance to rust isolates 12SD80 and 12NC29, was included as a positive experimental control for detection of H_2_O_2_. Three negative controls were used including an oat differential that contains the resistance gene *Pc14* which is not effective against 12SD80 and 12NC29, oat variety Marvelous infected with 12SD80 and 12NC29 and mock inoculated Marvelous. Samples were examined using an upright light microscope (Nikon Eclipse 90i). Mean and standard error of the mean of the number of urediniospores associated with H_2_O_2_ accumulation were calculated based on all three biological replicates. Digital micrographs were captured with a Nikon D2-Fi2 color camera using bright field and objective lens 4x and 10x. Multiple images were acquired in X, Y and Z planes using Z-series process using the Nikon Elements software. Z-stacking images were treated with an extended depth of focus (EDF) function to focus all the planes.

### qPCR quantification of fungal DNA

Genomic DNA was extracted from infected and mock-treated secondary leaves of *B. distachyon* accessions ABR6 and Bd21 at 3, 7, and 12 dpi using DNeasy Plant Mini Kit (Qiagen). The relative abundance of fungal DNA was measured using the Femto™ Fungal DNA Quantification Kit (Zymo Research) based on quantification of ITS regions using ITS-specific primers and fungal DNA Standards provided by the manufacturer. PCR was conducted using a CFX96 Real-Time system (Bio-Rad) and thermal cycles were set for initial denaturation at 95°C for 10 min, 45 cycles of 95°C for 30 s, 50°C for 30 s, 60°C for 60 s, followed by a final extension cycle at 72°C for 7 min. The *Brachypodium GAPDH* gene was also quantified in the same DNA samples using gene-specific primers (Hong et al. 2008). Fungal DNA level was normalized relative to the plant *GAPDH* value in each sample. Four independent experiments (biological replicates) were analyzed per time point per isolate, with two technical replicates within each biological replicate. Mean and standard error of the mean per condition (fungal isolate x *Brachypodium* accession x time point) were calculated based on data from all four experiments. Correlation between accumulation of fungal DNA, combined for all isolates at each time point (3, 7, and 12 dpi) and estimates of fungal colonization, combined for all isolates at 14 dpi was determined using a Pearson’s correlation coefficient test.

### RNA extraction and RT-qPCR analysis

Total RNA was extracted from infected and mock-treated secondary leaves of *B. distachyon* accessions ABR6 and Bd21 at 12, 24, 48, and 72 hpi using the RNeasy Plant Mini Kit (Qiagen). For RT-qPCR, cDNA was synthesized using the PrimeScript First Strand cDNA Synthesis Kit (Takara) and amplification was performed using the SensiFAST SYBR Lo-ROX Kit (Bioline). Sequences of gene-specific primers, reported by Gill et al. (2015) and Mandadi and Scholthof (2012) and those used in our study are listed in Supplementary Table 1. The *GAPDH, Ubi4* and *Ubi18* genes were examined as potential reference genes (Hong et al. 2008). Expression levels were normalized using the plant *GAPDH* gene after comparing PCR efficiencies and variation of quantification cycle (Cq) values for all three genes across time points in mock or pathogen-inoculated *Brachypodium* accessions (ABR6 and Bd21). PCR thermal cycles were set for initial denaturation at 95°C for 2 min, 40 cycles of 95°C for 5 s, followed by annealing/extension at 60°C for 20 s in a CFX96 Real-Time PCR system (Bio-Rad). Data was collected from three independent experiments (biological replicates) and each biological replicate included two technical replicates. Differential expression (DE) values were calculated as normalized fold changes of the expression using the ΔΔCT method (Livak and Schmittgen 2001), and DE value ≥ 2 was considered as a significant change in gene expression.

## Results

### Variation in resistance response of *Brachypodium* accessions to *P. coronata* f. sp. *avenae* infection

To examine the ability of *P. coronata* f. sp. *avenae* to infect non-host species *B. distachyon* and *B. hybridum* we evaluated the disease responses of 25 different *Brachypodium* accessions to three North American *P. coronata* f. sp. *avenae* isolates, 12SD80, 203, and 12NC29. First we characterized the virulence phenotypes of these isolates using a set of oat differentials (Fig. 1A). All three isolates are fully virulent on the susceptible oat variety Marvelous with obvious sporulation occurring at 14 dpi (Fig 1B, C). In contrast, no pustules were observed on any *Brachypodium* accession 30 days post-infection, which suggests that the asexual phase of the *P. coronata* f. sp. *avenae* life cycle cannot be completed on either *B. distachyon* or *B. hybridum*. However, macroscopic symptoms were observed on *Brachypodium* accessions in response to challenge with these rust isolates (Fig. 1D, E, Table 1) with some accessions developing chlorosis and/or necrosis of varying severity (Fig. 1F, Table 1). These symptoms were dependent upon both the accession genotype and rust genotype. For example, the necrosis severity of accession Bd1-1 was high in response to isolate 12SD80 but low upon infection with isolates 203 and 12NC29, whereas chlorosis severity in accession Adi13 was higher when challenged with isolates 12SD80 and 12NC29 but lower in response to isolate 203. Some *Brachypodium* accessions, such as Bd2-3, Bd3-1 and Tek4, reacted differently to all three rust isolates with different necrosis and chlorosis severities observed in each interaction. ABR6 consistently showed minimal symptom development, whereas accessions such as Bd21, Bd21-3 and Koz5 displayed more extreme necrosis and chlorosis responses to all three isolates.

**Fig. 1.**
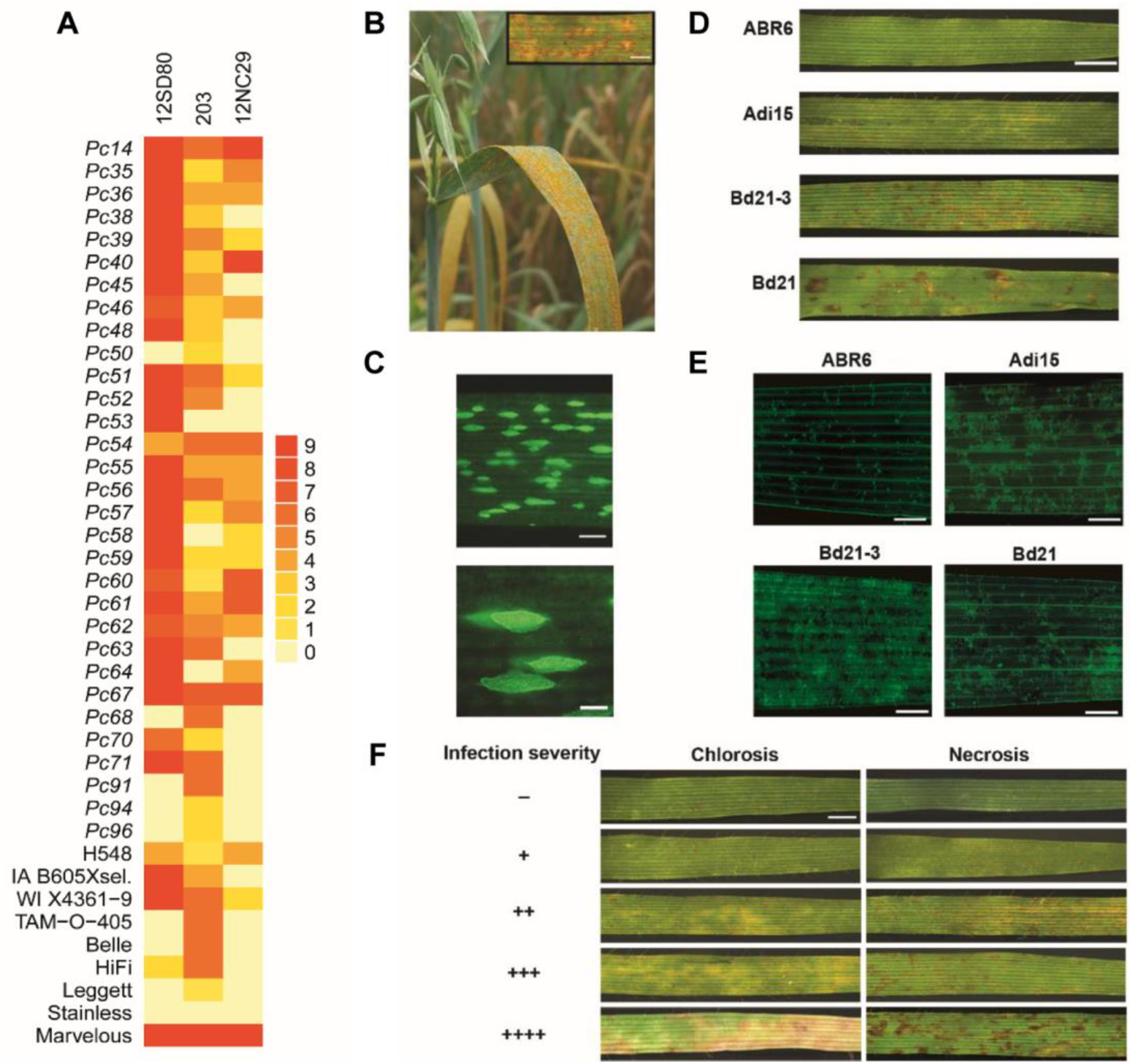
Infection of oat and *Brachypodium* accessions with *P. coronata* f. sp. *avenae*. **A**. Heat map of virulence phenotypes of *P. coronata* f. sp. *avenae* isolates on oat differentials. **B, C**. Formation of pustules and sporulation on infected susceptible oat leaves (Marvelous), respectively. **D, E**. Variation of infection symptoms and fungal colonization, respectively, on four *Brachypodium* accessions. **F**. Absence (−) or presence (+) of chlorosis and/or necrosis in *Brachypodium* accessions inoculated with *P. coronata* f. sp. *avenae* at 14 dpi. Level of symptom severity is indicated by the number of “+” characters. Symptoms correspond to the representative isolate 12SD80, except for the absence of necrosis which corresponds to isolate 203. Scale bars indicate 2 mm (B, D, F), 0.5 mm (C, top), 0.25 mm (C, bottom) and 0.5 mm (E)

**Table 1.** Variation of symptoms in *Brachypodium* accessions infected with *P. coronata* f. sp. *avenae* isolates.

| Accessions | 12SD80 |  | 203 |  | 12NC29 |  |
| --- | --- | --- | --- | --- | --- | --- |
|  | Chlorosis | Necrosis | Chlorosis | Necrosis | Chlorosis | Necrosis |
| ABR6 | - | + | - | - | - | + |
| ABR7 | + | + | + | + | + | + |
| Adi12 | +++ | + | ++ | + | + | + |
| Adi13 | +++ | ++ | + | +++ | ++++ | ++ |
| Adi15 | ++ | ++ | +++ | + | +++ | ++ |
| Bd1-1 | +++ | ++++ | +++ | + | + | ++ |
| Bd18-1 | - | + | + | + | +++ | ++ |
| Bd21 | ++ | +++ | +++ | +++ | ++ | ++ |
| Bd21-3 | ++ | +++ | +++ | ++ | ++++ | +++ |
| Bd2-3 | +++ | +++ | + | ++ | + | + |
| Bd29-1 | ++ | ++++ | + | + | + | +++ |
| Bd30-1 | + | ++ | + | + | + | +++ |
| Bd3-1 | ++ | ++++ | + | +++ | +++ | ++ |
| BdTR10H | ++ | ++ | + | + | ++ | ++ |
| BdTR13K | + | ++ | + | ++ | + | ++ |
| Foz1 | + | + | +++ | ++++ | + | - |
| J6.2 | + | + | + | + | + | + |
| Jer1 | + | +++ | - | + | + | ++ |
| Koz5 | ++ | ++ | +++ | ++ | +++ | ++ |
| Luc1 | + | + | + | - | ++ | + |
| Mon3 | ++ | ++++ | + | + | +++ | + |
| Tek4 | - | +++ | + | - | + | + |
| *Bel1 | + | ++ | + | + | + | ++++ |
| *Bou1 | + | + | + | ++ | - | +++ |
| *Pob1 | - | +++ | + | + | + | + |

Accessions with asterisks correspond to *B. hybridum* and those without asterisk are *B. distachyon* accessions. Symbols “+” and “-” show presence or absence of macroscopic symptoms, respectively, and number of “+” indicates symptom severity with ++++ as maximum level.

In parallel, all accessions were analyzed microscopically to determine the extent of *P. coronata* f. sp. *avenae* infection for each isolate. For all three isolates, urediniospores germinated to produce appressoria and penetrated the plant indicating that *Brachypodium* accessions are recognized as a potential host by *P. coronata* f. sp. *avenae* (Supplementary Fig. 1). A large variation in rust growth was observed amongst these *Brachypodium* accessions (Fig. 2A) although growth was always less than that observed on the oat host. Microscopically no *P. coronata* f. sp. *avenae* isolate showed initiation of uredinia on any *Brachypodium* accession confirming the non-host status of *B. distachyon* and *B. hybridum* to this phytopathogen.

**Fig. 2.**
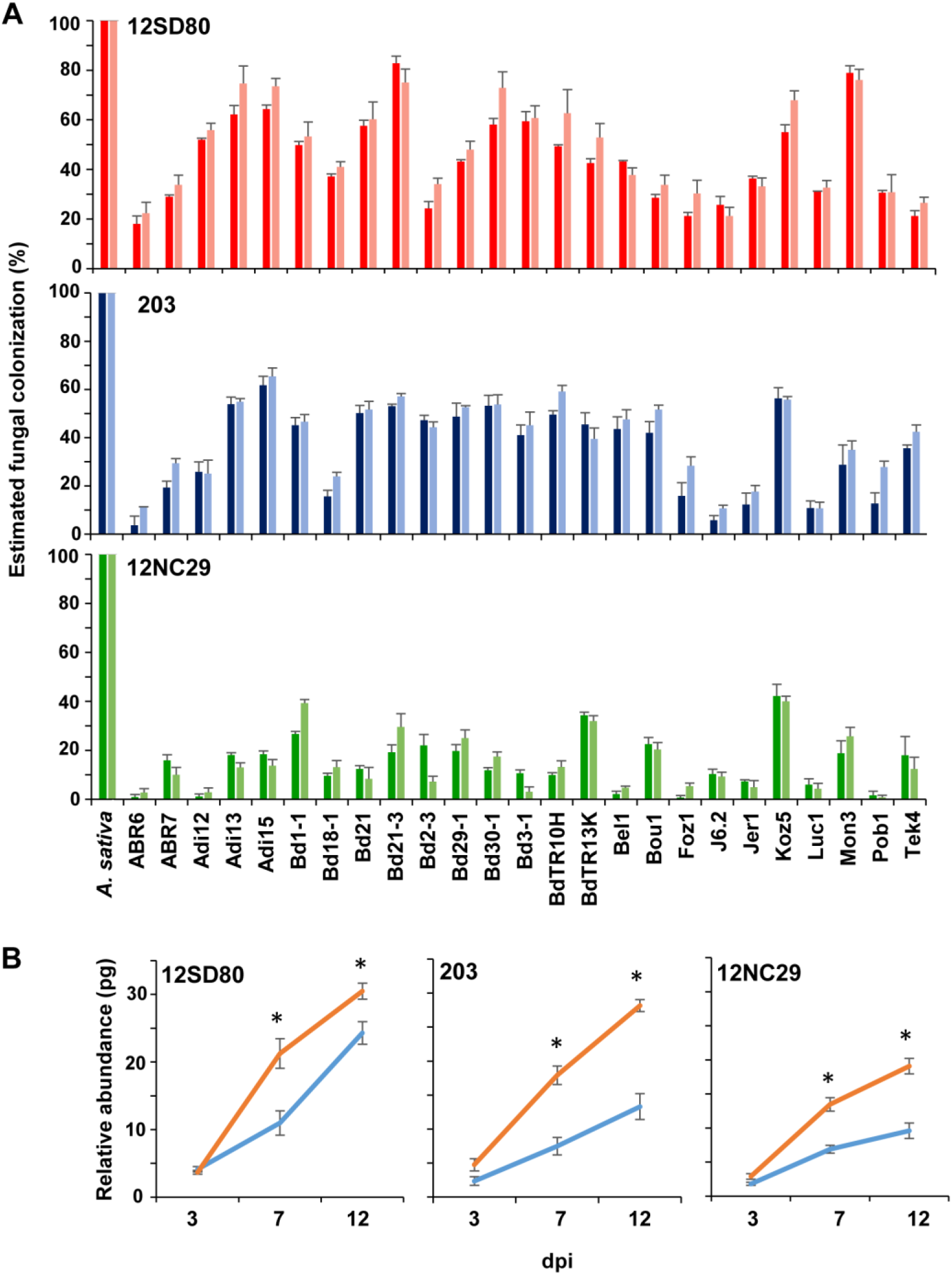
Colonization of *Brachypodium* accessions by *P. coronata* f. sp. *avenae*. **A**. Fungal growth estimates in foliar tissue for isolates 12SD80, 203, and 12NC29 depicted by the percentage of colonized area at 14 dpi. Crown rust susceptible oat (*A. sativa*) variety Marvelous serves as positive experimental control. Results from two independent experiments for each isolate are shown with distinct color intensities and each bar represents a mean value in one independent experiment (biological replicate). Error bars represent standard error of mean of three leaves (technical replicates) within one independent experiment. **B**. Quantification of fungal DNA for each rust isolate in *B. distachyon* accessions ABR6 (blue line) and Bd21 (orange line). Error bars represent standard error of mean of four independent experiments (biological replicates).

In general, *P. coronata* f. sp. *avenae* isolates 12SD80 and 203, which are more broadly virulent on the oat differentials (27 and 12, respectively) than isolate 12NC29 (5), showed more extensive growth on all tested *Brachypodium* accessions (Fig. 2A). Maximum colonization was greatest for isolate 12SD80, which colonized ∼79% of the leaf area of accession Bd21-3, while isolate 203 colonized ∼64% of the leaf area of accession Adi15. In contrast, the maximum colonization for isolate 12NC29 was ∼41% of the leaf area of accession Koz5. For all three isolates, Koz5 and Bd21-3 were amongst those *B. distachyon* accessions with the greatest fungal growth, whereas *B. distachyon* accessions ABR6 and Foz1, as well as *B. hybridum* accession Pob1, had the least fungal growth.

The fungal growth of each isolate and accession combination was scored relative to the most fungal growth observed for each isolate (Supplementary Fig. 2). Interestingly, some accessions (i.e., Bd30-1, Bd3-1, BdTR10H) were substantially infected by isolates 12SD80 and 203 while growth of 12NC29 was more restricted. In contrast, accession Tek4 was similarly infected by isolates 12SD80 and 12NC29, but had greater growth of 203, while isolate 12SD80 grew more prolifically on accession Mon3 than did either isolates 203 and 12NC29. Remarkably, we did not observe a correlation between the degree of necrosis and fungal colonization levels for each isolate (rho= 0.39-0.44, *p*=0.03-0.05, Spearman’s test). Weak correlations were found between chlorosis and colonization levels of 12SD80 (rho= 0.62, *p*=1.37e-06) and 203 (rho= 0.53, *p*=0.0064), in contrast to 12NC29 (rho= 0.30, *p*=0.15). For example, Bd29-1 showed extensive necrosis and chlorosis when inoculated with isolate 12SD80 in contrast to 203, but had high levels of colonization by both isolates. Accession Bd18-1 developed the highest necrosis and chlorosis in response to 12NC29 but supported the lowest growth of this isolate. In contrast, the most obvious symptoms on Jer1 correlated with the greatest fungal growth, in this case by isolate 12SD80.

The variation in fungal growth occurring on different *Brachypodium* accessions suggests that it may be possible to dissect the genetic architecture controlling this NHR against *P. coronata* f. sp. *avenae*. Given the availability of an ABR6 x Bd21 F_4:5_ *B. distachyon* family (Bettgenhaeuser et al. 2017) and the differences observed in *P. coronata* f. sp. *avenae* infection patterns (Fig. 2A), these two parental accessions were further analyzed. To further confirm the colonization values estimated for ABR6 and Bd21, fungal DNA accumulation was quantified by qPCR for each rust isolate over a time course experiment of 3, 7 and 12 dpi (Fig. 2B). Fungal DNA levels of each isolate increased in both accessions; however, statistical significant differences in the accumulation of DNA were observed between accessions at 7 and 12 dpi. Fungal DNA abundance was higher in accession Bd21 for all three isolates compared with ABR6, while isolate 12NC29 had the least growth on both accessions. A strong correlation was observed between levels of fungal colonization of all three isolates in ABR6 and Bd21 and their fungal DNA abundance at 7 dpi (r=0.93, *p*=0.0056, Pearson’s test) and 12 dpi (r=0.91, *p*=0.0098), while the correlation was not significant at 3 dpi (r=0.76, *p*=0.07). These results validate fungal colonization estimates shown in Fig. 2A.

### Development of infection structures of *P. coronata* f. sp. *avenae* isolates on *B. distachyon* ABR6 and Bd21

The development of infection structures (Fig. 3A) of isolates 12SD80, 203, and 12NC29 in a time course experiment (1, 2, 3, and 6 dpi) was compared between ABR6 and Bd21, with the susceptible oat variety Marvelous included as a positive control. Spore germination rates (GS) and appressorium development (AP) of all three rust isolates were similar on ABR6, Bd21 and oat (Fig. 3B). However, fungal growth on both *Brachypodium* accessions was significantly slower than on susceptible oat, with fewer substomatal vesicles (SV), haustorium-mother cells (HMC) and established colonies (EC) produced at 1, 2 and 3 dpi. The infection progression was generally slower on accession ABR6 than Bd21, while isolate 12SD80 grew at a faster rate on *B. distachyon* compared with the other two isolates, as demonstrated by the increased percentage of infection sites that formed SV in both accessions at 1 dpi. The most noticeable differences in the parental accessions ABR6 and Bd21 were related to the development of SV at 2 dpi with isolate 12SD80 and HMC at 3 dpi for isolates 12SD80 and 203.

**Fig. 3.**
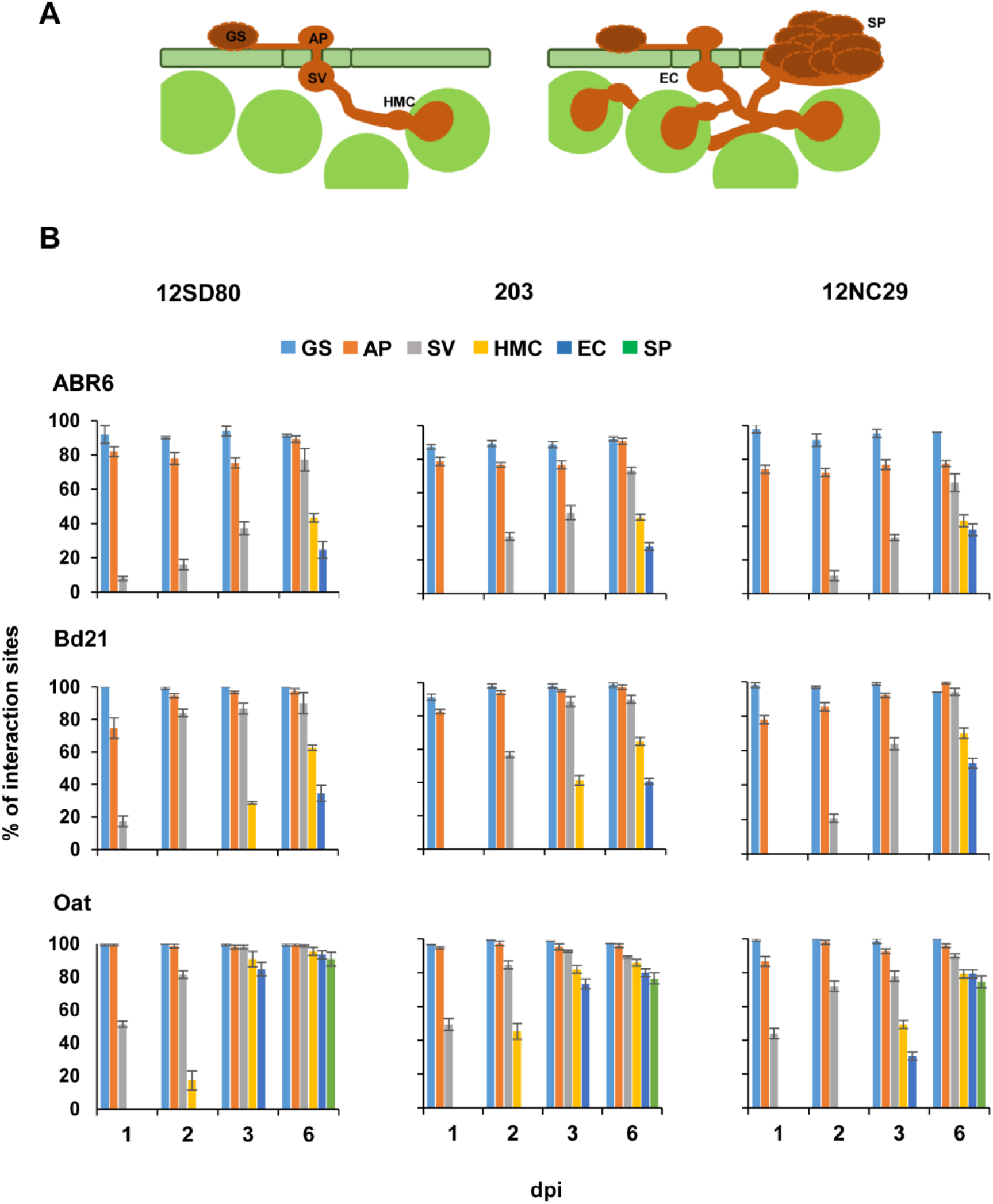
Fungal development of *P. coronata* f. sp. *avenae* in *Brachypodium* accessions. **A**. Illustration of *P. coronata* f. sp. *avenae* development in the plant. Germinated urediniospores (GS) form a penetration structure appressorium (AP) over a stoma. The fungus enters the mesophyll cavity and differentiates a substomatal vesicle (SV) and infection hypha. The establishment of a rust colony (EC) begins with the formation of a feeding structure (haustorium) which requires differentiation of a haustorial mother cell (HMC) near the hyphal tip. To undergo HMC formation, the rust fungus must come in contact with a mesophyll cell. **B**. Bars show the percentage of interaction sites with germinated urediniospores (GS, light blue), formation of appressorium (AP, orange), substomatal vesicle (SV, gray), haustorium-mother cell (HMC, yellow), established colony (EC, dark blue), and EC with sporulation (SP, green) in a sample of 100 infection sites per independent experiment (biological replicate). Error bars represent standard error of mean of three independent biological replicates.

### Histological analysis of reactive oxygen species in *B. distachyon* ABR6 and Bd21

The accumulation of H_2_O_2_ in mesophyll cells was examined by DAB staining in accessions ABR6 and Bd21 upon infection with isolates 12SD80, 203, and 12NC29 at 2, 4, and 6 dpi (Table 2, Supplementary Fig. 3). We conducted three independent experiments examining in each case 100 urediniospores from all three rust isolates. For isolate 12SD80 we found only a small number of urediniospores within 500 μm distance to DAB sites in ABR6 and Bd21, with slightly higher numbers of DAB sites in ABR6 than in Bd21 at 2 dpi and 4 dpi (Table 2). In contrast, DAB accumulation was rare in response to isolates 203 and 12NC29. We used an oat line carrying the *Pc91* gene, which confers resistance against 12SD80 and 12NC29 isolates (Fig. 1A), as a positive experimental control. In this case we observed a significantly greater proportion of urediniospores (∼50-60%) from 12SD80 and 12NC29 in association with H_2_O_2_ accumulation sites. The staining intensity and size of the H_2_O_2_ accumulation sites were also greater in the positive control samples than those in ABR6 and Bd21 (Supplementary Fig. 3). The resistance gene *Pc14*, which is not effective against 12SD80 and 12NC29 isolates, susceptible oat Marvelous infected with 12SD80 and 12NC29 (Fig. 1A) and mock inoculated Marvelous were used as negative controls and did not result in accumulation of H_2_O_2_ (Table 2, Supplementary Fig. 3). These data suggest that H_2_O_2_ accumulation is not a common feature of *B. distachyon* NHR to *P. coronata* f. sp. *avenae*.

**Table 2.** Histological analysis of H_2_O_2_ accumulation in *B. distachyon* accessions ABR6 and Bd21 and oat lines that contain the *Pc91* and Pc*14* genes in response to *P. coronata* f. sp. *avenae* infection.

| Isolates | Plants | 2 dpi | 4 dpi | 6 dpi |
| --- | --- | --- | --- | --- |
| 12SD80 | ABR6 | 10.7 ± 1.8 | 8.7 ± 2 | 4.3 ± 1.2 |
|  | Bd21 | 4 ± 1.5 | 3.3 ± 0.9 | 3.7 ± 1.5 |
|  | * Susceptible oat (Marvelous) | 0 | 0 | 0 |
| 203 | ABR6 | 0 | 0.3 ± 0.3 | 1.7 ± 0.3 |
|  | Bd21 | 0 | 0 | 2 ± 0.6 |
|  | * Susceptible oat (Marvelous) | 0 | 0 | 0 |
| 12NC29 | ABR6 | 0 | 0.7 ± 0.3 | 1 ± 0.6 |
|  | Bd21 | 0.3 ± 0.3 | 0 | 1 ± 0.6 |
|  | * Susceptible oat (Marvelous) | 0 | 0 | 0 |
| 12SD80 | ** Oat (*Pc91*) | 61.3 ± 7.5 | 65 ± 4.9 | 47.7 ± 4.1 |
|  | * Oat (*Pc14*) | 1 ± 0.6 | 0.3 ± 0.3 | 0.7 ± 0.7 |
| 12NC29 | ** Oat (*Pc91*) | 56 ± 9.5 | 54 ± 6.4 | 49 ± 5.9 |
|  | * Oat (*Pc14*) | 0 | 0.7 ± 0.3 | 0.7 ± 0.3 |

Values are calculated as means of three independent experiments, ± indicates standard error of the mean. Asterisks \*, \*\* indicate negative and positive experimental controls, respectively.

### Transcript profiling of defense-related genes in *B. distachyon* ABR6 and Bd21

To investigate the role of phytohormone-dependent defense responses in *Brachypodium* upon *P. coronata* f. sp. *avenae* infection, the temporal expression profile of several defense-related genes involved in SA, ET, and JA signaling pathways, as well as genes involved in callose synthesis and the phenylpropanoid pathway, were examined. This analysis was undertaken for ABR6 and Bd21 during the early stages of infection (up to 3 dpi), when there were not significant differences in fungal growth among isolates in both accessions (Fig. 2B). Three possible references genes were evaluated for data normalization, *GAPDH, Ubi4* and *Ubi18* (Hong et al. 2008). qRT-PCR primers for the *GAPDH* gene showed the highest amplification efficiency (98.4%) when compared to primers for *Ubi4* (96%) and *Ubi18* (97.1%) (Supplementary Fig. 4A). The *GAPDH* gene was selected for data normalization as it displayed the least variation in expression (Cq values) when compared to *Ubi4* and *Ubi18* in mock and fungal infected tissues in both *B. distachyon* accessions and all time points together (Supplementary Fig. 4B). *GAPDH* displayed steady expression in mock and infected tissues across time points in each accession (Supplementary Fig. 4C) which further supported the suitability of this gene for data normalization. In general, expression of genes involved in the SA and ET signaling pathways were induced in ABR6 and Bd21 during the first 48 h post-infection; however, expression of JA biosynthesis genes was not altered (Fig. 4, Supplementary Fig. 5). Changes in gene expression were greater in ABR6 than Bd21 upon infection with isolates 12SD80 and 203, but the inverse was observed for isolate 12NC29. These findings suggest differences between ABR6 and Bd21 in the early signaling responses to *P. coronata* f. sp. *avenae* infection.

**Fig. 4.**
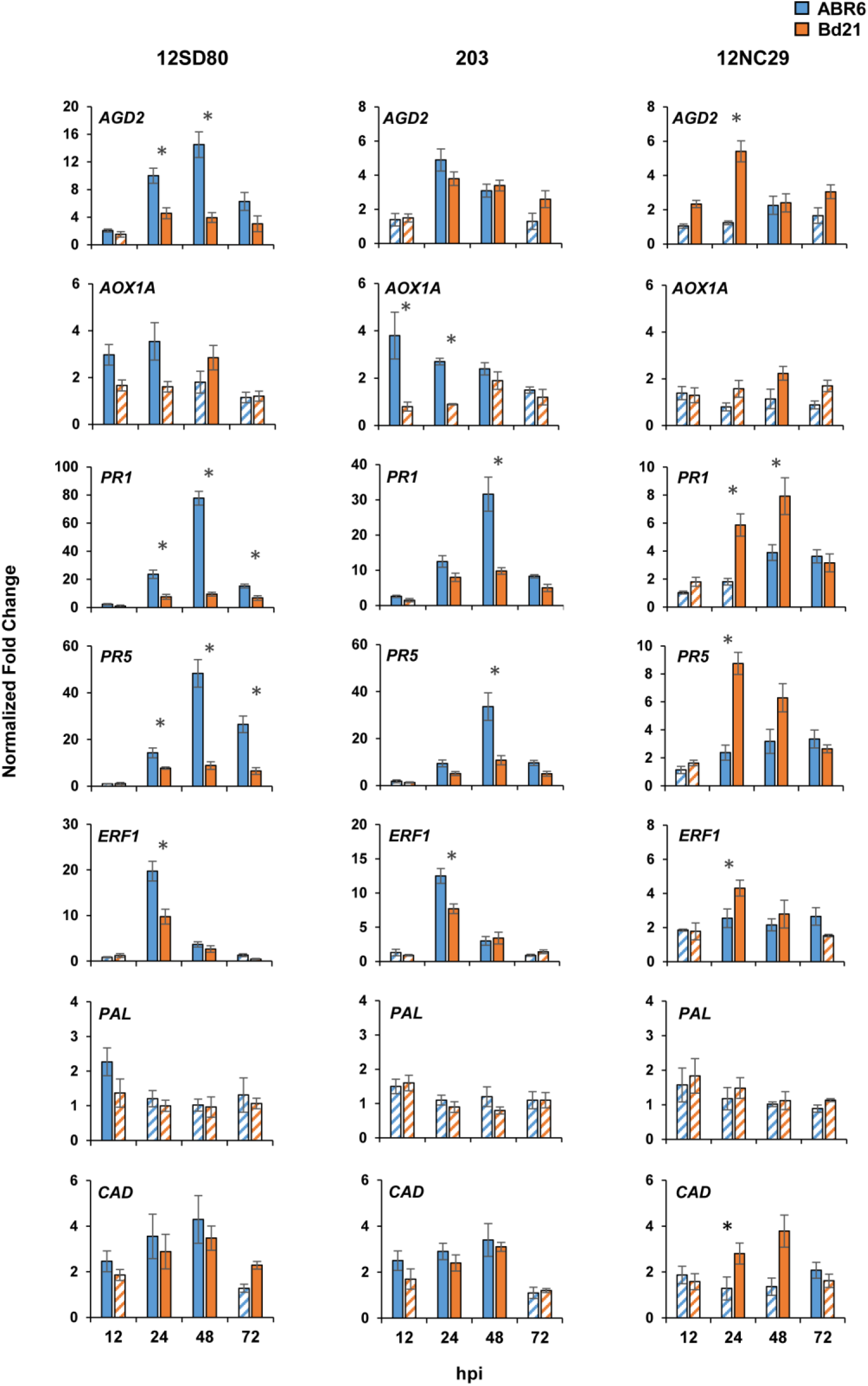
Expression profiling of various defense-related genes in *Brachypodium* accessions in response to *P. coronata* f. sp. *avenae*. Gene expression (fold-change) relative to mock inoculations in rust infected ABR6 (blue) and Bd21 (orange) plants of *Aberrant Growth Defects* 2 (*AGD2), Alternative Oxidase (AOX1A), Pathogenesis-related (PR) genes, Ethylene Response Factor* 1 (*ERF1*), *Phenylalanine Ammonia-Lyase* (*PAL*), and *Cinnamyl Alcohol Dehydrogenase* (*CAD*). Barplots represent mean values of fold change per time point. Solid colored bars indicate a fold change value of ≥ 2 whereas hatched bars indicate values below this threshold. Error bars represent standard error of mean of of three independent experiments (biological replicates). Asterisks indicate statistically significant differences (*p* ≤ 0.05) between ABR6 and Bd21 accessions as determined by a *t*-test.

Among the SA signaling pathway genes, expression of *Aberrant Growth Defects 2* (*AGD2*) peaked at 24 to 48 hpi with isolates 12SD80, 203 and 12NC29. The highest upregulation of *AGD2* was observed in the interaction of 12SD80 and ABR6. Expression of *Alternative Oxidase* (*AOX1A*) was induced in ABR6 as early as 12 hpi with isolates 12SD80 and 203, but it was not affected in response to isolate 12NC29. In contrast, *AOX1A* was upregulated in Bd21 only at 48 hpi in response to isolates 12SD80 and 12NC29. The *Pathogenesis-related* (*PR*) genes *PR1* and *PR5* showed the greatest induction amongst all genes tested in response to all three crown rust isolates. The expression of both genes peaked at 48 hpi except for *PR5* expression after inoculation with 12NC29 which peaked at 24 hpi. Overall, induction of these SA-responsive genes occurred in all interactions, but were remarkably stronger in ABR6 when inoculated with 12SD80 and 203, while Bd21 showed similar responses to all three isolates.

Expression of *Ethylene Response Factor 1* (*ERF1*) was maximal at 24 hpi with all three isolates and, as observed for SA-responsive genes, expression of *ERF1* was particularly high in ABR6 in response to isolates 12SD80 and 203. However, expression of an ET biosynthesis gene, *Aminocyclopropane-1-carboxylic Acid Oxidase* (*ACO1*), two JA biosynthesis genes, *Lipoxygenase 2* (*LOX2*), and *12-oxophytodienoate Reductase 3* (*OPR3*), *WRKY18* transcription factor, and *callose synthase* did not change in either of the accessions in response to challenge with any of the three isolates (Supplementary Fig. 5). Expression of *Phenylalanine Ammonia-Lyase* (*PAL*) was only slightly induced in ABR6 at 12 hpi with isolate 12SD80. *Cinnamyl Alcohol Dehydrogenase* (*CAD*) reached maximum induction in both accessions at 24 to 48 hpi with isolates 12SD80 and 203. In response to infection with isolate 12NC29, this gene was only upregulated in Bd21 between 24 to 48 hpi.

### Transcript profiling of *BdSTP13* in *B. distachyon* ABR6 and Bd21

The expression of the *Brachypodium* ortholog of *Lr67* (*STP13*) (Bradi1g69710) (Moore et al. 2015) was examined in accessions ABR6 and Bd21 upon infection with all three rust isolates, as it acts as a putative hexose transporter. *BdSTP13* was induced in both accessions in response to the tested rust isolates (Fig. 5). Gene induction peaked at 24 to 48 hpi, except in the interaction between ABR6 and12NC29, which showed low *BdSTP13* transcript accumulation.

**Fig. 5.**
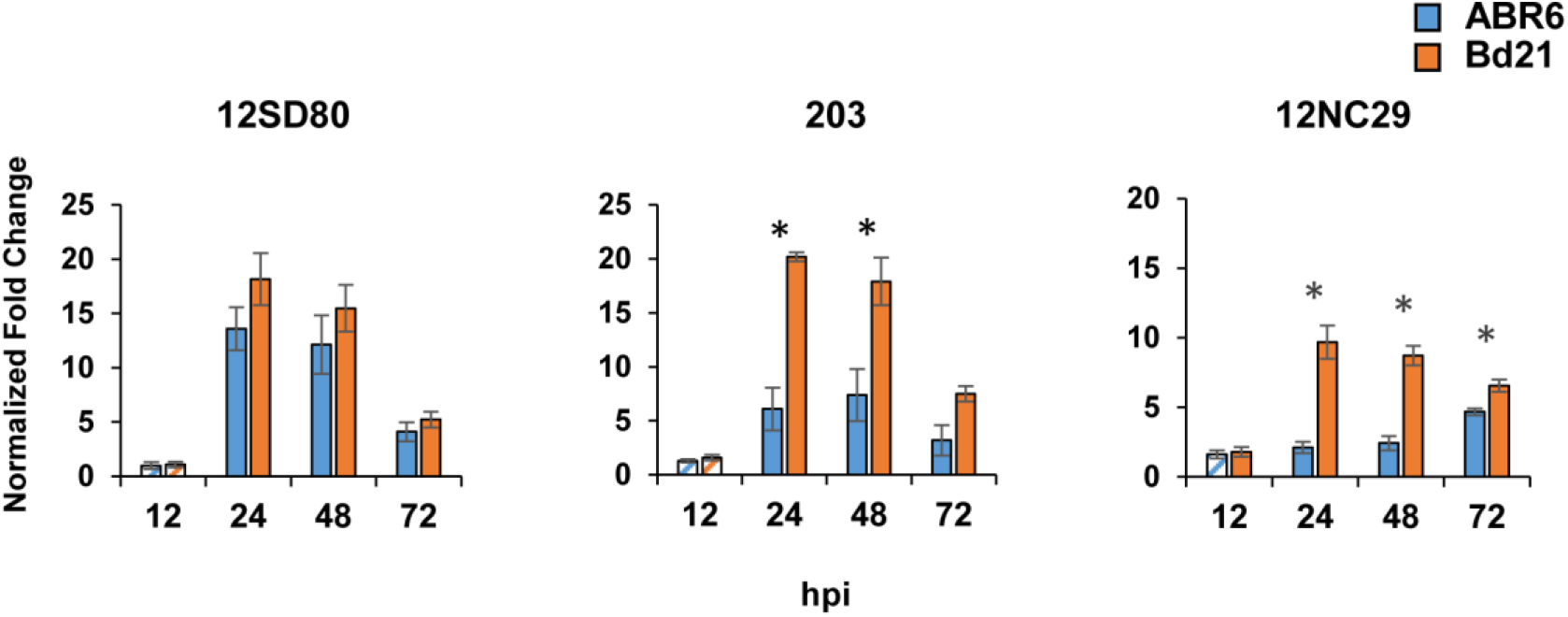
Expression profiling of *BdSTP13* in *Brachypodium* accessions in response to *P. coronata* f. sp. *avenae*. Gene expression (fold-change) relative to mock inoculations in rust infected ABR6 (blue) and Bd21 (orange) plants of the putative hexose transporter *BdSTP13*. Solid colored bars indicate a fold change value of ≥ 2 whereas hatched bars indicate values below this threshold. Error bars represent standard error of mean of three independent experiments (biological replicates) Asterisks indicate statistical significant differences (*p* ≤ 0.05) between ABR6 and Bd21 accessions as determined by a *t*-test.

## Discussion

The lack of effective *P. coronata* f. sp. *avenae* resistance in oat coupled with rapid evolution of pathogen virulence necessitates a search for new sources of resistance to control oat crown rust disease. To explore the potential of using *Brachypodium* species as a germplasm resource for disease resistance against *P. coronata* f. sp. *avenae*, we have characterized the interaction between three *P. coronata* f. sp. *avenae* isolates and a panel of *Brachypodium* accessions, including *B. distachyon* and *B. hybridum*. At the macroscopic level, variation in resistance phenotypes was observed, which has been previously reported for other rust interactions with *B. distachyon*. In these studies, *P. striiformis, P. graminis*, and *P. triticina* infection phenotypes varied from immunity to a range of symptoms that included pustule formation and sporulation (Ayliffe et al. 2013; Figueroa et al. 2013; Garvin 2011). However, similar to *Brachypodium-P. emaculata* (switchgrass rust) interactions (Gill et al. 2015), we did not observe *P. coronata* f. sp. *avenae* sporulation on any *Brachypodium* accession, which supports the status of *B. distachyon* and *B. hybridum* as non-hosts for this pathogen. Macroscopic infection symptoms were often associated with chlorosis and/or necrosis; however, it was visually difficult to confirm if these symptoms were a consequence of *P. coronata* f. sp. *avenae* infection. Microscopic analysis showed that *P. coronata* f. sp. *avenae* isolates 12SD80, 203, and 12NC29 overcame pre-haustorial resistance defenses and were able to colonize leaves of all *Brachypodium* accessions tested. These results are similar to the interactions observed between *P. graminis* f. sp. *tritici* and *B. distachyon* (Ayliffe et al. 2013). In light of these observations, we would like to emphasize that our definition of *B. distachyon* and *B. hybridum* as non-hosts is based on previous established terminology, which uses absence of sporulation as indicator of resistance. However, given that both species tolerate growth of *P. coronata* f. sp. *avenae*, both *B. distachyon* and *B. hybridum* could be considered as non-native hosts, and previous observations reflect also variations in susceptibility. Furthermore, changes in growth conditions may favor sporulation of *P. coronata* f. sp. *avenae* in both *B. distachyon* and *B. hybridum*.

To better understand the phenotypic variation existing between *P. coronata* f. sp. *avenae and Brachypodium* interactions, we compared the extent of fungal growth on a range of accessions. The rust isolates included in this study represent three distinct physiological races (genotypes). A wide range of pathogen growth was observed on different *Brachypodium* accessions and was dependent on both plant and fungal genotypes. Some *Brachypodium* accessions tolerated more fungal growth when infected with isolates 12SD80 and 203 versus 12NC29 (i.e., Bd30-1, Bd3-1, BdTR10H) (Fig. 2, Supplementary Fig. 2). Interestingly, we also identified accessions (i.e., Tek4) that tolerated more growth of isolates 12SD80 and 12NC29 than 203. These findings could be explained either by the lack of a plant target for pathogen effectors to promote infection effectively or the presence of race-specific components to resistance. The latter scenario implies that resistance is likely governed by variation in effector repertoires of the rust isolates and *R* genes present in the *Brachypodium* accessions. Given the close evolutionary relationship between oat and *Brachypodium* spp., it is possible that the accessions included in our study carry *R* genes that can detect effectors of *P. coronata* f. sp. *avenae*. Isolates 12SD80 and 12NC29 display different virulence profiles on the oat differential set and the recent sequencing of these isolates provides evidence of extensive differences in effector gene complements of 12SD80 and 12NC29 (Miller et al. 2018). Future studies identifying and comparing the effector repertoires of rust species, including those in our study and others that infect *Brachypodium* (i.e., *Puccinia brachypodii*) may help to explain our findings.

The precise contribution of ETI to non-host rust resistance in *Brachypodium* remains elusive. Future efforts to dissect the genetic factors controlling resistance to *P. coronata* f. sp. *avenae* in *B. distachyon* will help to evaluate the model proposed by Schulze-Lefert and Panstruga (2011), in which ETI is the major contributor to NHR in plant species that are closely related to the natural host. Heterologous expression of fungal and bacterial effectors in several non-host pathosystems supports that effector recognition by immunoreceptors contributes to resistance (Adlung et al. 2016; Giesbers et al. 2017; Lee et al. 2014; Stassen et al. 2013; Sumit et al. 2012), although in some cases effector recognition can occur in the absence of resistance (Giesbers et al. 2017; Goritschnig et al. 2012). Close examination of *Brachypodium* interactions with *P. graminis* ff. spp. *tritici, avena* and *phalaridis, P. triticina*, and *P. striiformis* suggests that HR-induced cell death is rare (Ayliffe et al. 2013). However, the lack of HR in these interactions does not necessarily undermine the contribution of ETI to non-host resistance, as there are instances when *R* gene-mediated ETI (e.g., wheat stem rust resistance gene, *Sr33*) can result in resistance without cell death (Periyannan et al. 2013). Further studies are needed to better understand the extent of ETI and HR contributions in NHR given that molecular and genetic factors in these interactions could be pathosystem-specific.

To investigate the role of ROS in *B. distachyon* response to *P. coronata* f. sp. *avenae*, we examined accumulation of ROS in accessions ABR6 and Bd21, which display contrasting infection phenotypes. We found a small number of urediniospores from 12SD80 associated with ROS accumulation, while detection of ROS was rare for other two isolates. However in these cases the accumulation of ROS in ABR6 and Bd21 was substantially less than that detected in a resistant oat line carrying the resistance gene *Pc91* (Table 2). These findings make the involvement of this ROS unlikely in the resistance of *Brachypodium* to *P. coronata* f. sp. *avenae*. In contrast to H_2_O_2_, phytohormones may play a thus far undefined role in modulating NHR in *Brachypodium* to rust pathogens.

Plant responses to pathogens are partly regulated through a complex interplay between salicylic acid (SA), ethylene (ET) and jasmonic acid (JA) signaling pathways (Denancé et al. 2013). Gill et al. (2015) found that several defense-related genes involved in SA, ET, and JA signaling pathways were induced in *Brachypodium* accessions infected with *P. emaculata*. In our study, upregulation of several SA-responsive genes and the key ethylene response regulator *ERF1* was detected, suggesting that SA and ET-signaling pathways may positively regulate NHR against *P. coronata* f. sp. *avenae*. The *Pathogenesis-related* (*PR*) genes *PR1* and *PR5* were upregulated in response to all rust isolates and given that *PR1* is a key marker gene of SA signaling (Kouzai et al. 2016; Sels et al. 2008), this further supports that the activation of SA-dependent defense responses may contribute to the phenotypes observed in *B. distachyon* and *B. hybridum* during *P. coronata* f. sp. *avenae* infection. JA biosynthesis genes were not induced, making this hormone unlikely to play an active role in this NHR response. In contrast, *ERF1*, which acts as a key regulatory element in the ET/JA-dependent defense responses, was upregulated suggesting that ET-responsive genes may enhance NHR (Berrocal-Lobo et al. 2002; Lorenzo et al. 2003; Müller and Munné-Bosch 2015). *Phenylalanine Ammonia-Lyase* (*PAL*) is a key enzyme in biosynthesis of polyphenolic compounds and lignin precursors and is often associated with host resistance responses against pathogens (Kalisz et al. 2015). Lignin deposition in response to pathogen attack is usually correlated with reinforcement of the cell wall and enhanced resistance (Miedes et al. 2014). Upregulation of *PAL* does not appear to play a significant role in the responses of *B. distachyon* to *P. coronata* f. sp. *avenae;* however, induction of *CAD* may point to cell wall alterations in response to isolates 12SD80 and 203. A genome-wide transcriptional analysis of the interaction of *P. coronata* f. sp. *avenae* with ABR6 and Bd21 will help to elucidate the involvement of some of these specific processes in NHR, including phytohormone regulation.

Extensive *P. coronata* f. sp. *avenae* growth in some *Brachypodium* accessions implies that the pathogen is capable of nutrient accession for a period of time and possibly able to target susceptibility factors conserved between *Brachypodium* and oat. Little is known about the mechanisms underlying rust susceptibility. Pathogens can alter sugar partitioning in the host to accommodate their growth (Lapin and Van den Ackerveken 2013), and thus sugar transporter proteins (STPs) may be targeted by rust fungi to increase nutrient availability (Dodds and Lagudah 2016). The wheat *STP13* hexose transporter is implicated in supporting rust pathogens as mutations in this gene leads to broad-spectrum resistance (Moore et al. 2015). We examined the expression of the ortholog of wheat *STP13* in *Brachypodium* during infection by *P. coronata* f. sp. *avenae*. Interestingly, the highest levels of *BdSTP13* expression were obtained in accession Bd21, which is subjected to more rust growth than ABR6, suggesting that the gene may indeed serve as a susceptibility factor. Our results are consistent with previous findings that show that expression of *Lr67*, and its homoeologs in wheat peaks at 24 hpi upon infection with the leaf rust fungus, *Puccinia triticina*. Gene expression analysis of *Lr67* as well as other *STP13* genes in *Arabidopsis* and grapevine, indicates that the gene is also induced in response to pathogens (Hayes et al. 2010; Lemonnier et al. 2014; Moore et al. 2015). These observations provide a basis to study the contribution of sugar transport to rust susceptibility in an experimental system that is not as complex as in hexaploid oat and wheat.

Several studies have demonstrated the value of mining wild relatives of crop species to introduce disease resistance. Effective race-specific rust resistance has been introgressed into wheat (*Triticum aestivum*) from related species, such as the *Sr50* gene from rye (*Secale cereale)*, which confers resistance to wheat stem rust by recognition of the corresponding AvrSr50 effector of *P. graminis* f. sp. *tritici* (Chen et al. 2017; Mago et al. 2015). Similarly, functional transfer of the pigeonpea (*Cajanus cajan*) *CcRpp1* resistance gene into soybean confers resistance against soybean rust (*Phakopsora pachyrhizi)*, which demonstrates how heterologous resistance transgenes can be used for crop improvement (Kawashima et al. 2016). This interspecies transfer of disease resistance highlights the possibility that *Brachypodium* genes that confer NHR to *P. coronata* f. sp. *avenae* could potentially be transferred to oat and provide disease resistance. As a next step, we are utilizing an ABR6 x Bd21 mapping population to identify resistance loci that could be tested in *A. sativa*. Both accumulation of fungal DNA and analysis of development of fungal infection structures (i.e., SV or HCM) can be used to phenotype the ABR6 x Bd21 mapping population. In summary, our findings indicate that *Brachypodium* is a suitable species for evaluating NHR to *P. coronata* f. sp. *avenae* and is a potential source of novel disease resistance for oat. The rapid expansion of genomic resources, including fully sequenced genomes and assessment of genetic diversity among *Brachypodium* genotypes (International *Brachypodium* Initiative 2010; Gordon et al. 2014, 2017) enables this species to be exploited for engineering disease resistance in closely related crop species.

## Acknowledgements

We acknowledge support by the University of Minnesota Experimental Station USDA-NIFA Hatch/Figueroa project MIN-22-058, as well as the USDA-ARS-The University of Minnesota Standard Cooperative Agreement (3002-11031-00053115) between S.F.K and M.F. M.J.M. is supported by the Gatsby Foundation and Biotechnology and Biological Sciences Research Council (BB/P012574/1). We thank P.N. Dodds and E. Henningsen for discussions and comments during manuscript preparation, as well as Roger Caspers and Lief van Lierop for technical support. We would like to acknowledge the University Imaging Centers (UIC) (http://uic.umn.edu) and staff support at the University of Minnesota for using microscopy equipment.

## Supplementary files

**Supplementary Fig. 1.**
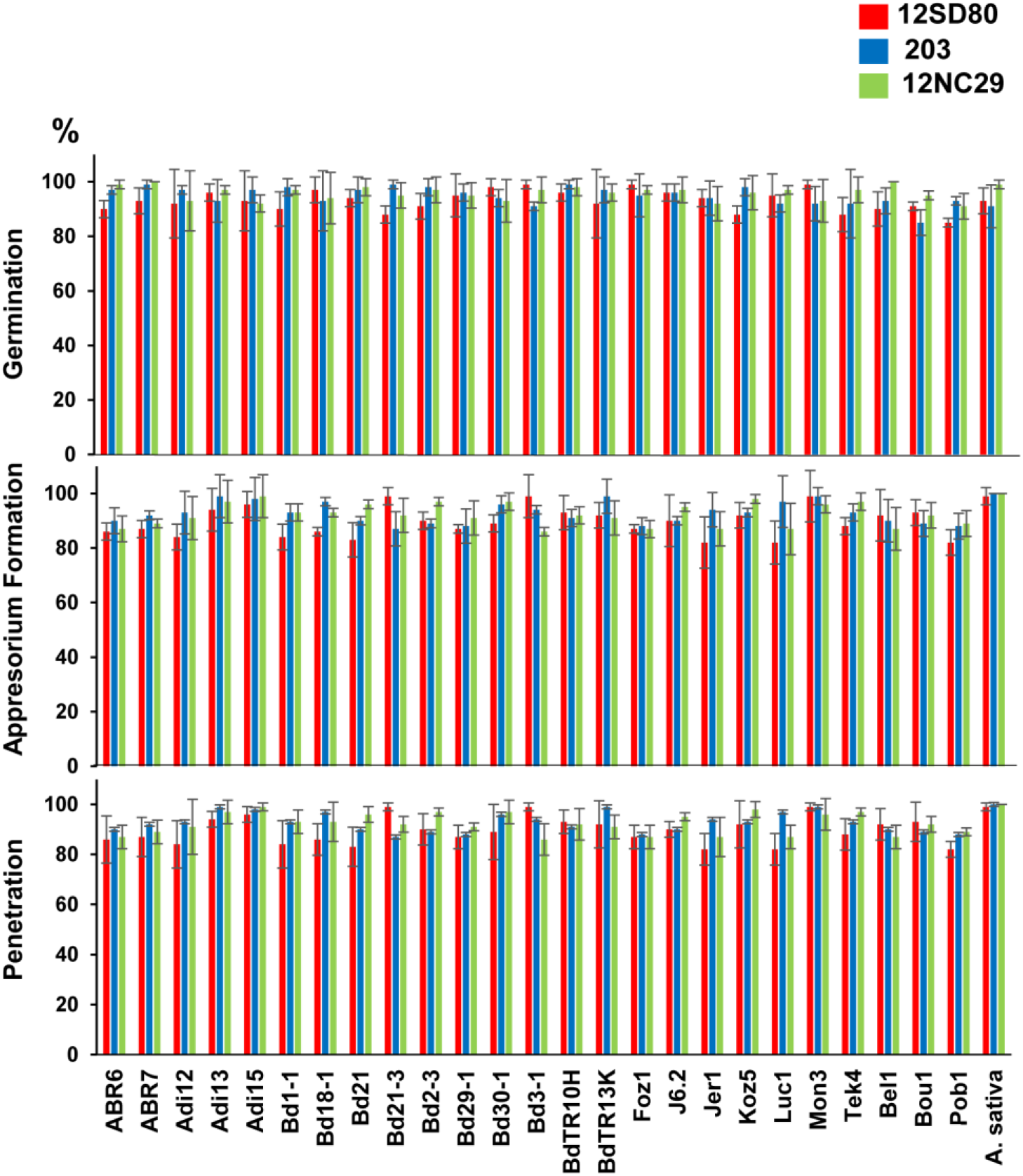
shows development of *P. coronata* f. sp. *avenae* in all *Brachypodium* accessions and susceptible oat Marvelous as quantified by percentage of germination, formation of appressorium and occurrence of plant penetration. Error bars represent standard error of mean of two independent experiments (biological replicates). Data was collected for 50 infection sites per independent biological replicate.

**Supplementary Fig. 2.**
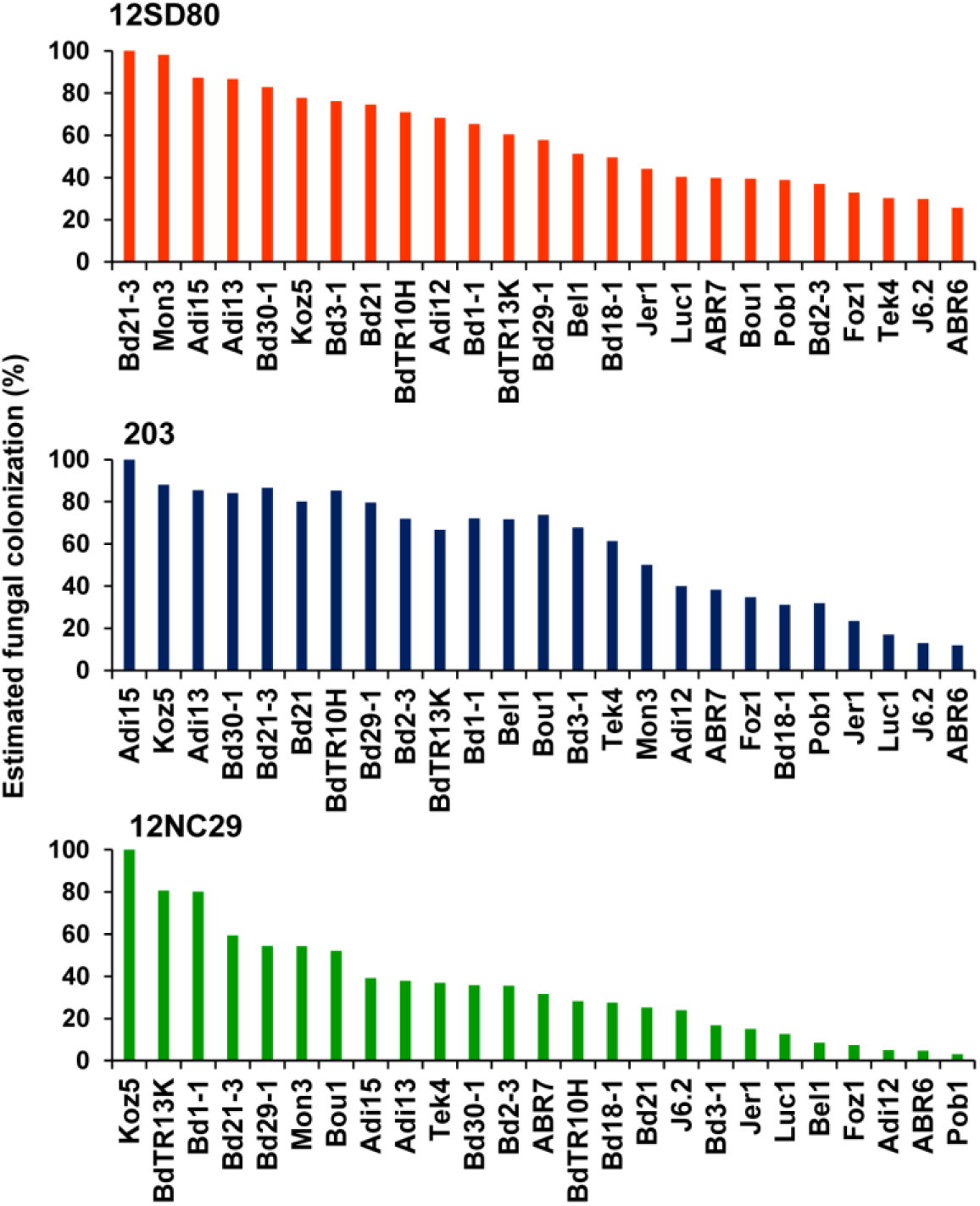
shows colonization of *Brachypodium* accessions by *P. coronata* f. sp. *avenae*. Fungal growth estimate per isolate, with individual isolates 12SD80, 203 and 12NC29 shown as a percentage value relative to the accession displaying the highest level of colonization for each isolate.

**Supplementary Fig. 3.**
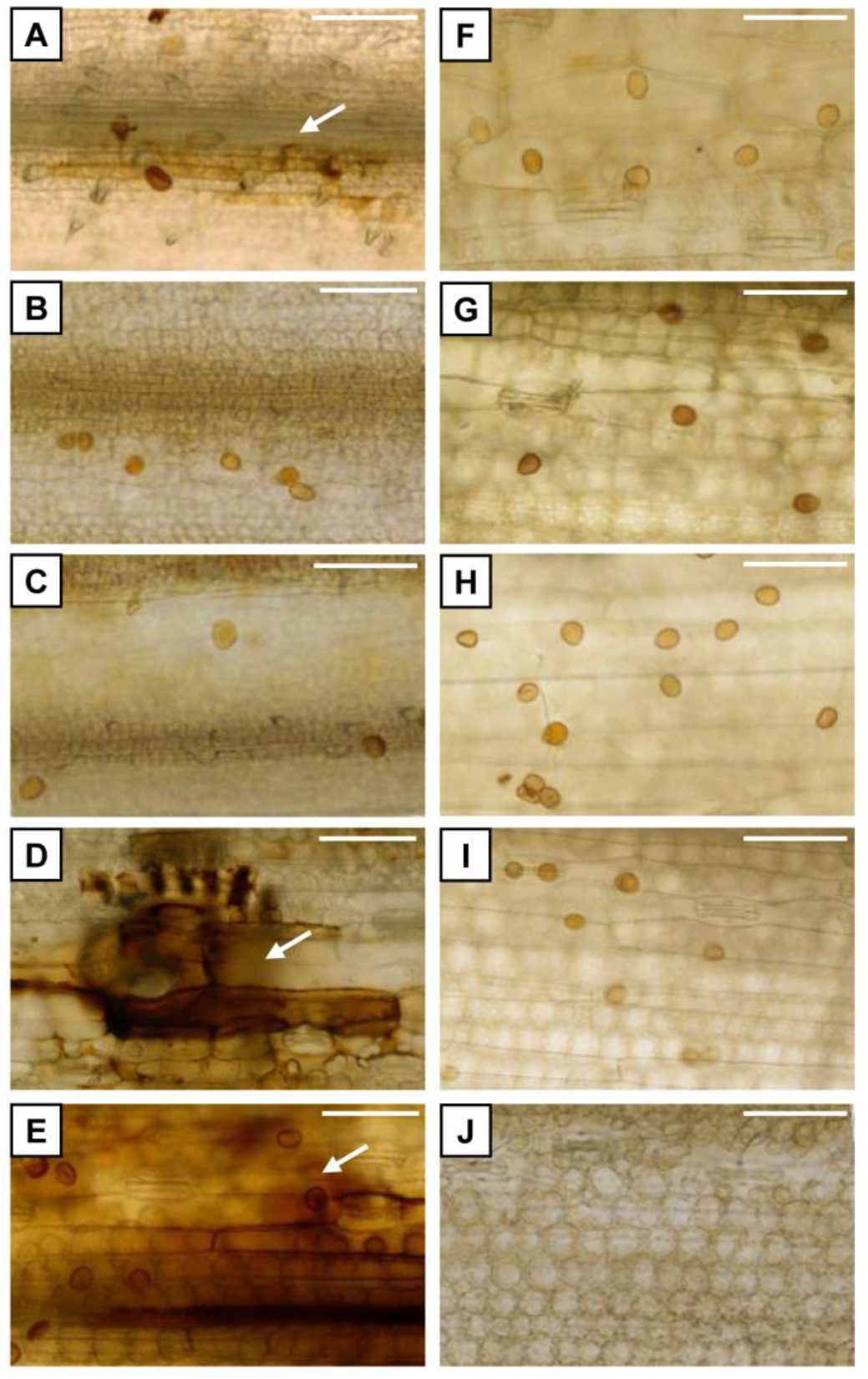
shows examples of H_2_O_2_ detection at 2 dpi using DAB staining. Images depict urediniospores and presence of H_2_O_2_ is shown by a white arrow. **A, B, C**. *B. distachyon* accession ABR6 inoculated with rust isolates 12SD80, 203, and 12NC29, respectively. **D, E**. Oat differential line that contains the resistance gene *Pc91* inoculated with isolates 12SD80 and 12NC29, respectively. The gene *Pc91* is effective against both isolates and therefore serves as positive experimental control. **F, G**. Oat differential line that contains the resistance gene *Pc14* inoculated with isolates 12SD80 and 12NC29, respectively. The gene *Pc14* is not effective against any of the isolate and therefore serves as negative experimental control. **H, I**. Susceptible oat variety Marvelous inoculated with isolates 12SD80 and 12NC29, respectively. **J**. Lack of H_2_O_2_ in mock inoculated oat Marvelous (negative control). Scale bars indicate 100 μm.

**Supplementary Fig. 4.**
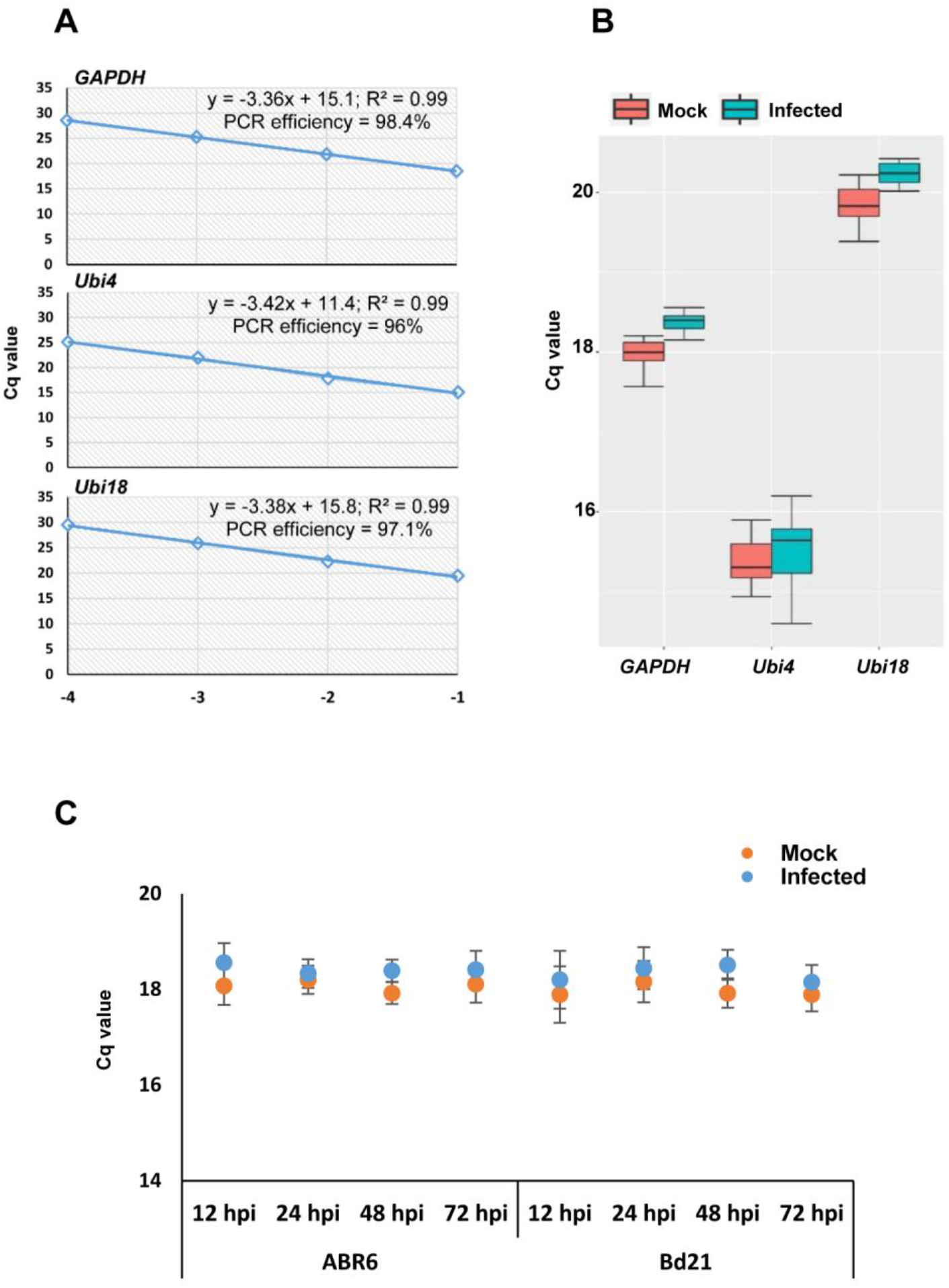
shows evaluation of *GAPDH, Ubi4* and *Ubi18* as potential reference genes for data normalization in targeted gene expression analyses via qRT-PCR. **A**. Amplification efficiency of primers for *GAPDH, Ubi4* and *Ubi18* genes. **B**. Comparison of expression levels of *GAPDH, Ubi4*, and *Ubi18* genes between mock and pathogen-infected tissues after combining Cq values from both *B. distachyon* accessions (ABR6 and Bd21) and all time points (12, 24, 48, and 72 hpi). Boxplots show variation between mean Cq values and whiskers on boxplots show variability outside the upper and lower quartiles. Data was collected from three independent experiments each including three technical replications. **C**. Comparison of expression levels of *GAPDH* in ABR6 and Bd21 in both mock and infected samples across time points. Graph shows mean values and error bars on dotplots represent standard error of mean of three independent experiments (biological replicates).

**Supplementary Fig. 5.**
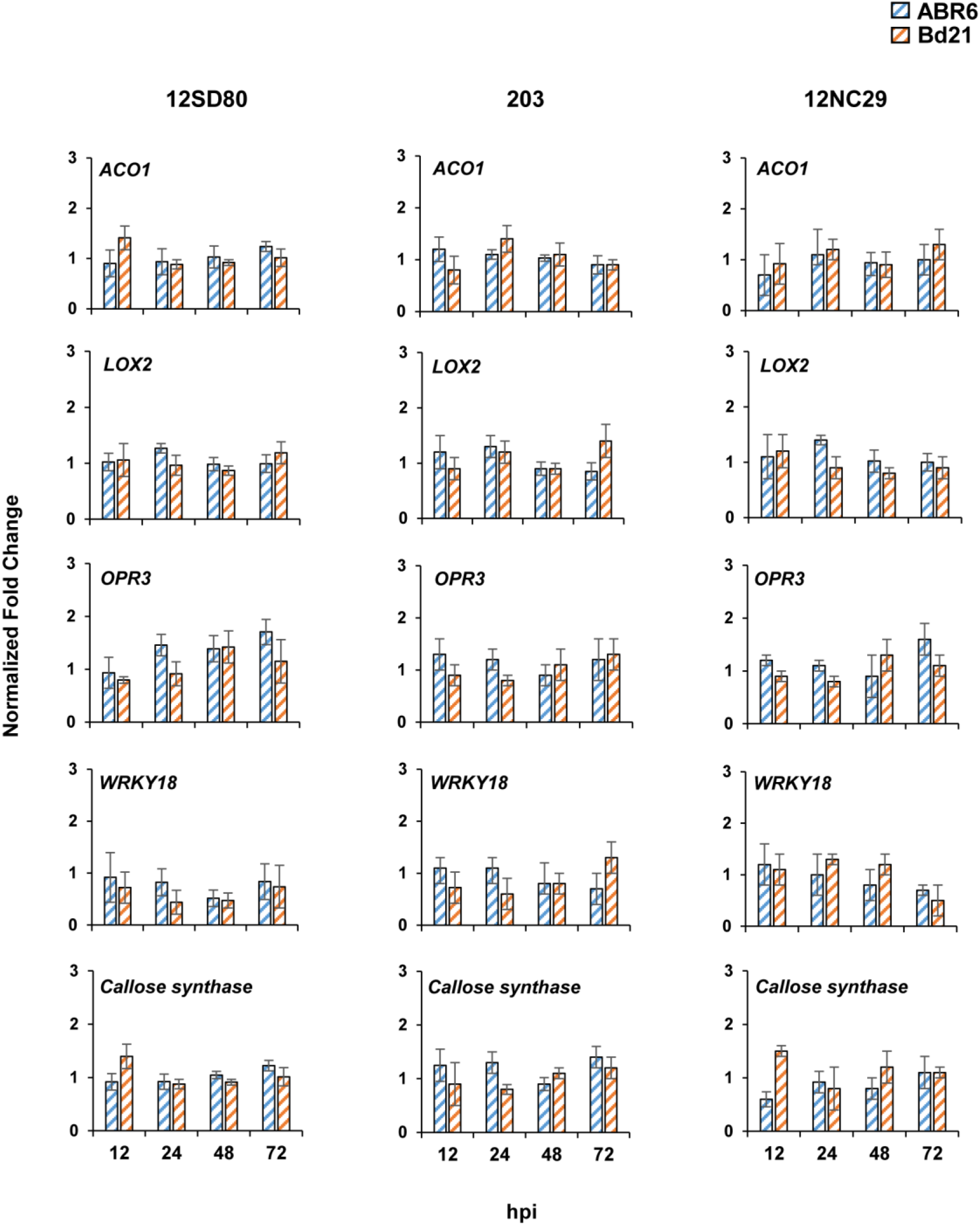
shows expression profiling of various defense-related genes in *B. distachyon* accessions in response to *P. coronata* f. sp. *avenae*. Gene expression (fold-change) relative to mock inoculations in rust infected ABR6 (blue) and Bd21 (orange) plants of *Aminocyclopropane-1-carboxylic Acid Oxidase (ACO1), Lipoxygenase 2* (*LOX2*), *12-oxophytodienoate Reductase 3* (*OPR3*), *WRKY18* transcription factor, and *callose synthase*. Barplots represent mean values of fold change per time point. Solid colored bars indicate a fold change value of ≥ 2 whereas hatched bars indicate values below this threshold. Error bars represent standard error of mean of three independent experiments (biological replicates). Asterisks indicate statistical significant differences (*p* ≤ 0.05) between ABR6 and Bd21 accessions as determined by a *t*-test.

**Supplementary Table 1.**
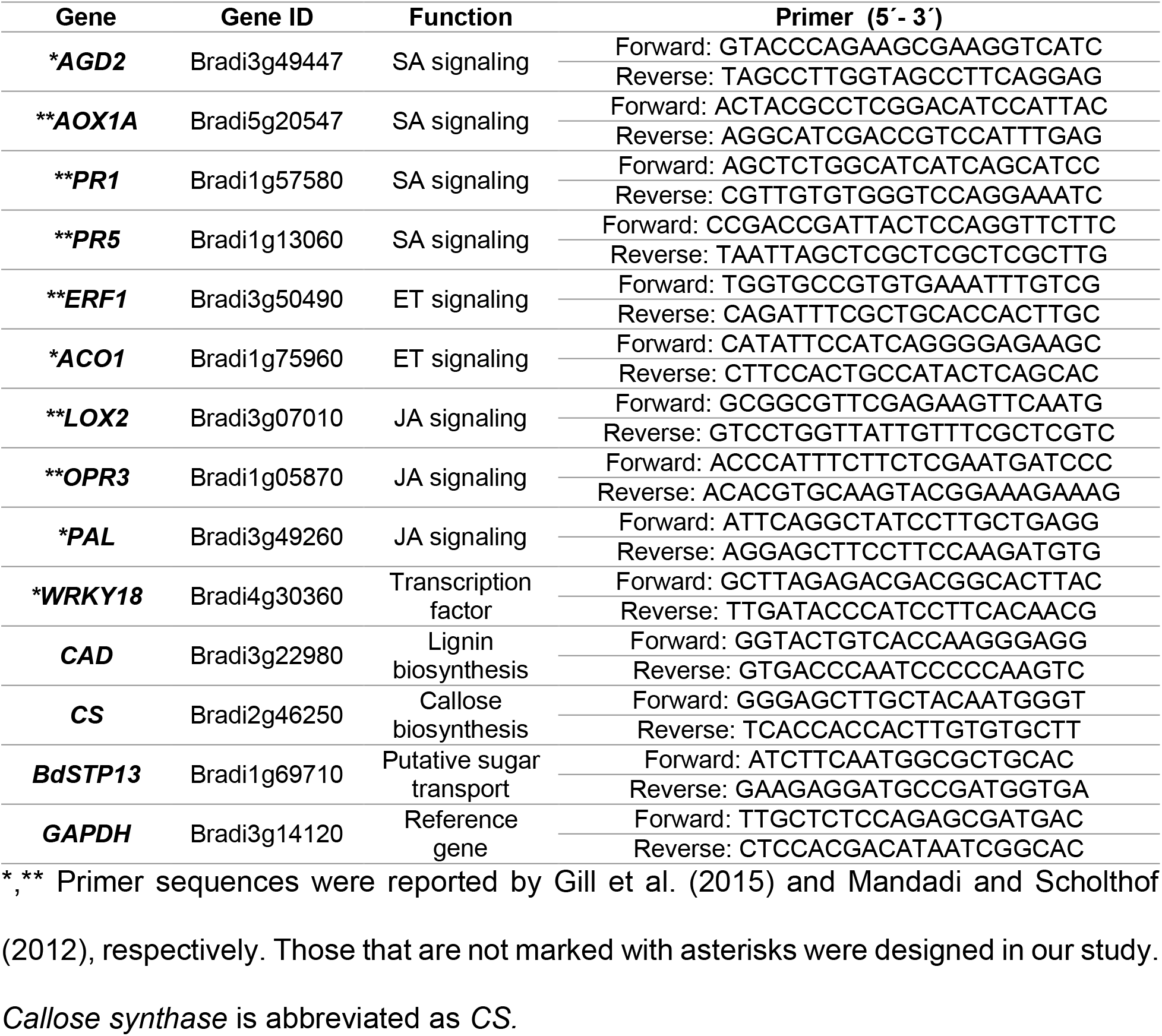
List of gene-specific primers used for RT-qPCR analysis.

